# Polar opposites; bacterioplankton susceptibility and mycoplankton resistance to ocean acidification

**DOI:** 10.1101/2020.02.03.933325

**Authors:** Storme Zaviar de Scally, Samuel Chaffron, Thulani Peter Makhalanyane

**Author notes:** Correspondence to: Thulani P. Makhalanyane, Centre for Microbial Ecology and Genomics (CMEG), Department of Biochemistry, Genetics and Microbiology, University of Pretoria, Hatfield, Pretoria, 0028, South Africa.

## Abstract

Microorganisms form the basis of ocean ecosystems yet the effects of perturbations such as decreasing pH on microbial community structure, interactions and functionality remain compared to multicellular organisms. Using an experimental manipulation of Southern Ocean seawater, we subjected bacterioplankton and mycoplankton to artificial pH decreases, which are predicted to occur in the future. We show that acidification led to substantial increases of bacterioplankton diversity, while in contrast it had no effect on mycoplankton diversity. Our analyses revealed a loss of putative keystone taxa and a decrease in predicted community interactions as a response to lower pH levels. Bacterioplankton shifted from generalist to specialist community members, suggesting a specific stress response to unfavourable conditions. In addition, enzyme activities involved in nitrogen acquisition were lower at reduced pH levels, suggesting altered organic matter cycling in a more acidic ocean. Our findings suggest that bacterioplankton and mycoplankton may respond differentially to future ocean acidification, with potentially negative impacts on community structure and biogeochemical cycling in the Southern Ocean.

**IMPORTANCE:** Oceans absorb the majority of anthropogenically produced CO_2_, the consequence of which is ocean acidification, a phenomenon already negatively impacting key marine organisms. Marine microbial communities form the basis of ocean food webs by generating nutrients for higher trophic levels, yet the response of these key microbial drivers to acidification remains unclear. This knowledge deficit is particularly true for understudied marine ecosystems such as the Southern Ocean. Using a mesocosm approach, we found that acidification severely impacts microbial community stability, by altering bacterioplankton community structure, reducing network complexity, and augmenting enzyme activities associated with nitrogen acquisition. This study adds to our understanding of the effects of ocean acidification on microbial communities, particularly within an environment expected to be largely effected by future anthropogenically driven climate change.

## INTRODUCTION

The ocean absorbs roughly 30% of anthropogenic carbon dioxide emissions, which leads to changes in oceanic carbonate chemistry, resulting in both carbonation and acidification (Orr et al., 2005; Zeebe et al., 2008; Doney et al., 2009; Adler et al., 2011; Kottmeier et al., 2016). Globally, ocean pH is predicted to decrease by as much as 0.4 and 0.8 units within the next 100 and 300 years, respectively (Caldeira and Wickett, 2003; Orr et al., 2005), a change which is nearly eight times higher than what has been observed within the last 25 million years (Pearson and Palmer, 2000). Ocean acidification (OA) can negatively impact marine macro-organisms, such as calcifiers, adversely affecting their growth and physiology (Doney et al., 2009; Gazeau et al., 2013; Brandenburg et al., 2019). However, the extent to which decreasing pH affects diverse bacterioplankton and mycoplankton communities remains largely unclear.

A null hypothesis proposed by Joint and colleagues (2011b) states that due to the ubiquitous abundance, fast generation time and genetic plasticity of microorganisms, microorganisms will rapidly adapt to future pH perturbations. However, despite the large genetic flexibility microorganisms possess, a physiological tipping point, where individual organisms are no longer able to adapt to perturbations, may be reached (Allison and Martiny, 2008b; Hutchins and Fu, 2017). There is considerable uncertainty and disagreement regarding the impact of acidification on bacterioplankton and mycoplankton communities, with reports demonstrating both resistance and susceptibility. Evidence of microbial community resistance to acidification include: unchanged bacterial cell abundance (Grossart et al., 2006; Allgaier et al., 2008; Krause et al., 2012; Newbold et al., 2012), consistent composition (Lindh et al., 2013; Roy et al., 2013; Oliver et al., 2014), unaffected or increased heterotrophic production (Grossart et al., 2006; Arnosti et al., 2011) and increased nitrogen fixation (Joint et al., 2011a; Rees et al., 2016). Susceptibility has been seen through, alterations in bacterial and fungal cell numbers (Krause et al., 2013; Endres et al., 2014; Engel et al., 2014), compositional shifts (Allgaier et al., 2008; Krause et al., 2012; Zhang et al., 2013) with changes towards pathogenic members (Vega-Thurber et al., 2009; Reich et al., 2017) and decreased nitrification rates (Beman et al., 2011). Moreover, the increased expression of proton pump genes identified in a metatranscriptomic study (Bunse et al., 2016) suggests that acidification may be energetically costly for microbial cells, with possible negative implications for microbial functionality and biogeochemical cycling. These results emphasise the need for further investigations into the effects of acidification on complex and diverse bacterioplankton and mycoplankton assemblages (Hutchins and Fu, 2017).

Although the Southern Ocean (SO) covers less than 20% of the global ocean it is a highly productive ecosystem, facilitating the inter-ocean exchange of nutrients which drives over 70% of ocean productivity (Mayewski et al., 2009). While the SO has naturally low pH, it remains vulnerable to anthropogenic induced acidification due to its’ low temperature, the upwelling of CO_2_ enriched deep waters, and the fact that approximately 40% of the global oceanic uptake of CO_2_ occurs in the SO (Riebesell and Gattuso, 2015). Several studies have shown that acidification results in increased resilience in diatoms (Valenzuela et al., 2018) and functional biodiversity loss (Teixidó et al., 2018). In contrast to other environments (Newbold et al., 2012; Chen et al., 2013; Huggett et al., 2018; James et al., 2019), the precise responses of bacterioplankton and mycoplankton remain largely speculative and often unexamined in pristine marine environments. This knowledge is crucial for understanding the effects of climate change in regions most sensitive to future environmental change.

To address this deficit, we assessed the effect of future predicted pH levels (based on IPCC projections for the years 2100 and 2300, respectively) on both SO bacterioplankton and mycoplankton diversity, composition, predicted interactions, and functionality using mesocosm experiments. We predicted that; (1) ocean acidification will decrease both bacterioplankton and mycoplankton diversity and will significantly alter community structure; (2) microbial functionality will be both positively and negatively affected by ocean acidification in terms of carbon and nitrogen/phosphorus decomposition, respectively, and (3) the total number of covariations and subsequently network stability will decrease with lower pH levels.

## RESULTS

### Microbial diversity and community structure in control and acidified mesocosms

Alpha diversity for bacteria and archaea was significantly higher in the low and medium pH groups in comparison to the control (ANOVA, p-value < 0.001), and both Shannon diversity (Fig.1a) and OTU richness (Fig.1b) significantly correlated with pH. On the contrary, fungal alpha diversity was not significantly different between the three treatment groups (ANOVA, p-value > 0.05), and did not correlate significantly with pH (Fig. 1ab). Bacterial and archaeal community structure was significantly affected by pH treatment (PERMANOVA, p-value < 0.05, betadisper p-value > 0.05), and clustered according to treatment type (Fig. 2a). In contrast to bacteria and archaea, fungal community structure was not significantly influenced by pH treatment (PERMANOVA, p-value < 0.05, betadisper p-value < 0.05), as no distinct clusters were seen according to treatment group (Fig. 2b). Thus, bacterioplankton and mycoplankton community composition clearly display differential responses to acidification. Whereas several bacterioplankton genera either increased or decreased in abundance under acidified conditions, mycoplankton composition remained stable, with the exception of Agariomyctes. Specific genera and species differences are further described in Supporting Information (Fig. S3).

**Fig. 1:**
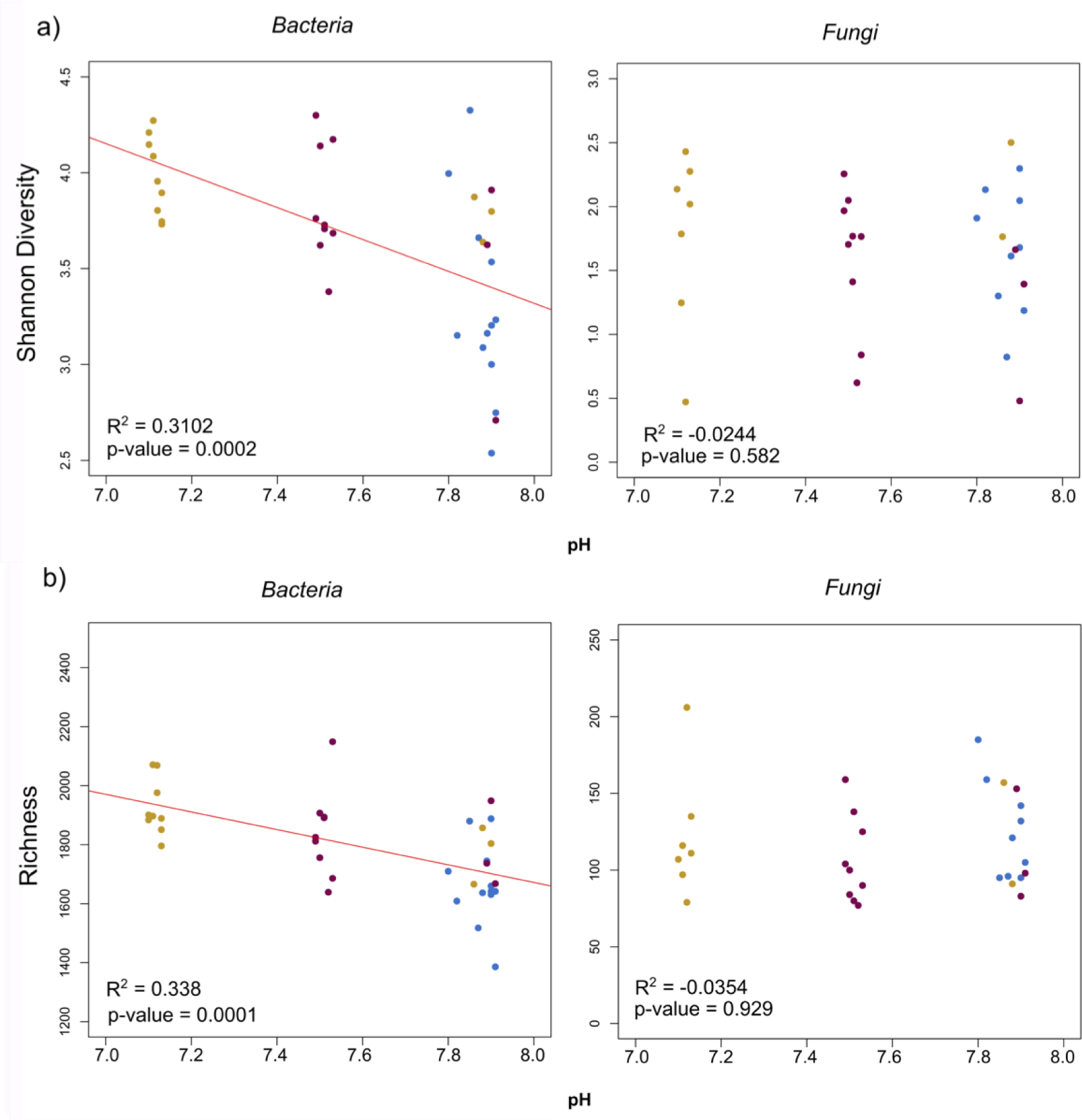
Relationships between bacterial and fungal diversity and experimentally induced seawater pH changes. Shannon diversity or H’ index (a) and richness or observed species (b) are indicated in the top and bottom panels, respectively. Solid lines represent the fitted regression model. The percentage of variance explained by the model or R^2^, as well as the significance level of the model, is indicated in the bottom left corner of each sub-figure. Fungal diversity relationships with pH were non-significant (p-value > 0.5), and therefore no regression line is shown.

**Fig. 2:**
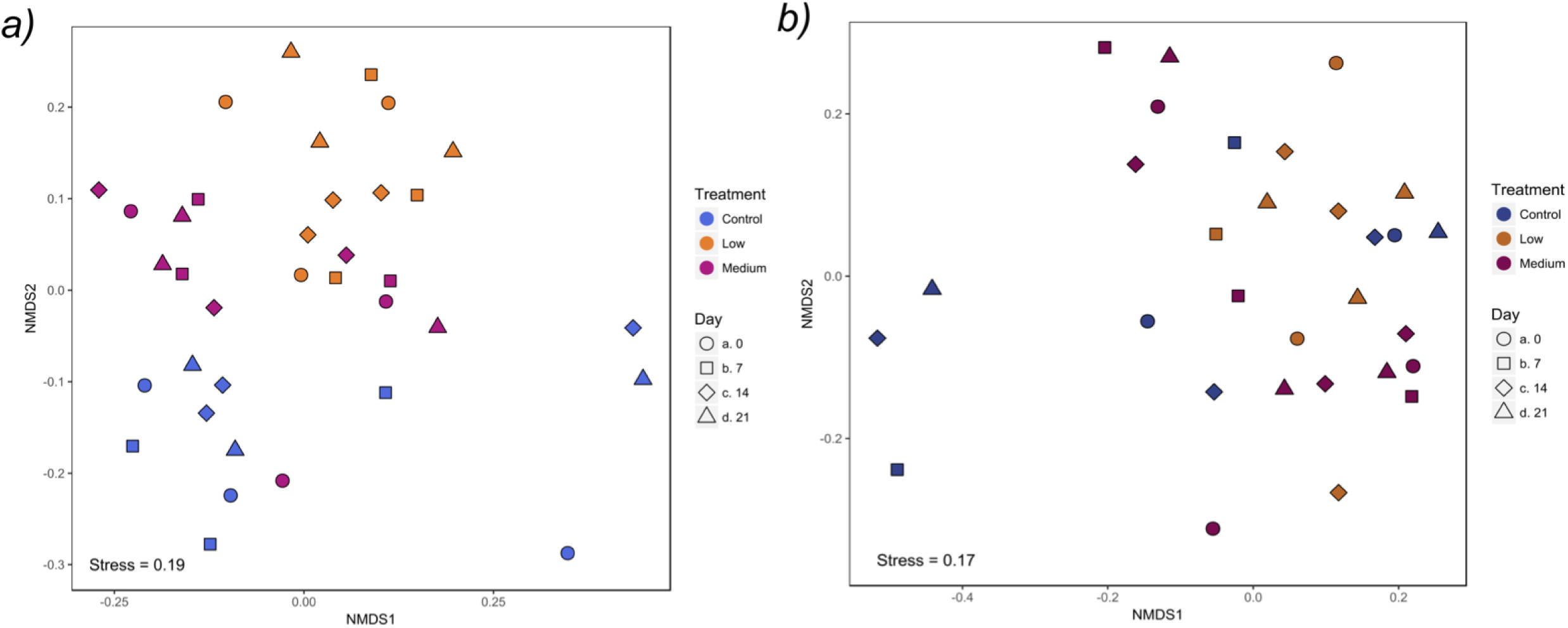
Non-metric multi-dimensional scaling (nMDS) ordination plots of bacterioplankton and mycoplankton communities constructed using Bray-Curtis dissimilarity. The left (a) and right (b) panels represent the 16S rRNA gene and ITS 1 region datasets, respectively. Points represent samples and are coloured and shaped according to treatment group and sampling day, respectively. Stress values are indicated at the bottom of each sub-figure.

### Functional community responses to ocean acidification

Extracellular enzymatic activities were detected in all samples, where the activities of carbon and phosphorus acquiring enzymes did not vary significantly with pH treatment (Fig. 3). In contrast, nitrogen acquiring enzymes NAG and LAP significantly decreased at lower pH. Enzymes involved in carbon acquisition, namely BG and BX, showed no significant increase in response to pH treatment (ANOVA, p-value > 0.05). AP, involved in phosphorus acquisition, showed no significant increase or decrease in activity in response to the pH treatment (ANOVA, p-value > 0.05). NAG activity decreased in the low group across sampling days, although this was not significant from the control group (ANOVA, p-value > 0.05). Notably, LAP was shown to be significantly lower in the medium and low pH treatment groups relative to the control at day 21 of the mesocosm experiment (ANOVA, p-value < 0.05). Activities were correlated to pH levels using both linear regression (Fig. 3) and Spearman’s rho. Both results indicated that only NAG was significantly positively associated with pH (*r* = 0.42, p-value < 0.05; Fig. 3a), whereby NAG activity decreased at lower pH levels. LAP activity, although significantly different between treatment and control groups, could not be predicted by pH, likely due to the lower activity in the medium pH group than the low pH group.

**Fig. 3:**
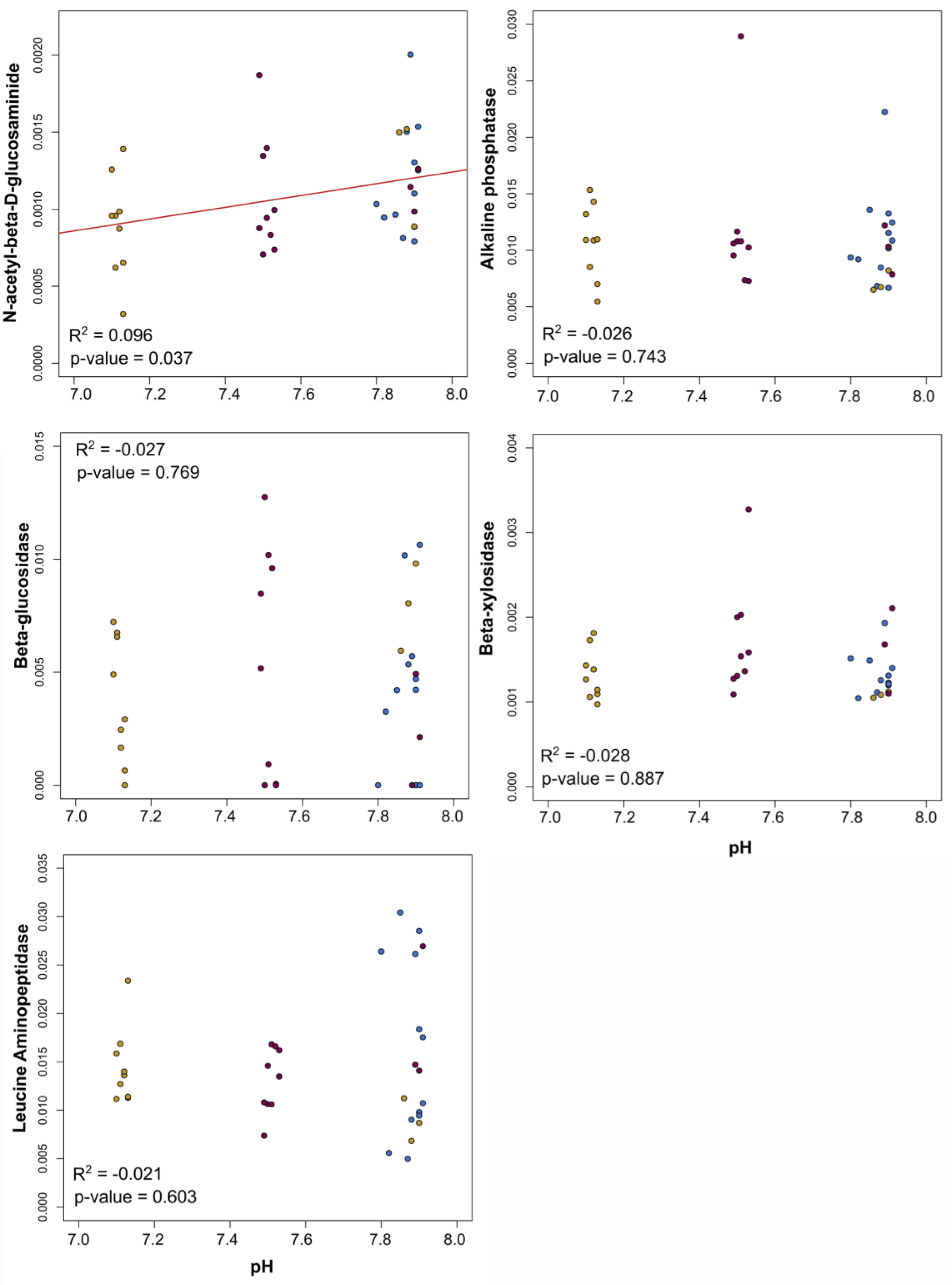
Relationship between extracellular enzymatic activities and experimentally induced acidification. The solid red line represents the fitted regression model. The percentage of variance explained by the model or R^2^, as well as the significance level of the model, is indicated in the left corner of each sub-figure. No fitted regression line is shown for all exoenzyme activities which were not significantly correlated with pH (p-value > 0.5). Points are coloured by the three treatment groups where orange, purple and blue indicate low, medium and control, respectively.

### Community structures and system level changes associated to ocean acidification

Table S2 shows topological features of bacterial and archaeal community networks. All networks exhibited scale-free characteristics, typical of biological association networks, were small world, modular, and significantly different from generated random networks (see Supporting information for details).

Network structure, complexity and community composition differed markedly between the three treatment groups (Table S2, Fig. 4). The medium network was the largest followed by the low and control networks, showing that acidification resulted in larger networks. Interestingly, although the medium network displayed the highest diversity and connectivity, the low network was the least complex (Table S2). The medium network contained the greatest number of edges, the largest average degree (i.e. average number of edges per node in the network) and the shortest harmonic geodesic distance (i.e. distance between all nodes in the network).

**Fig. 4:**
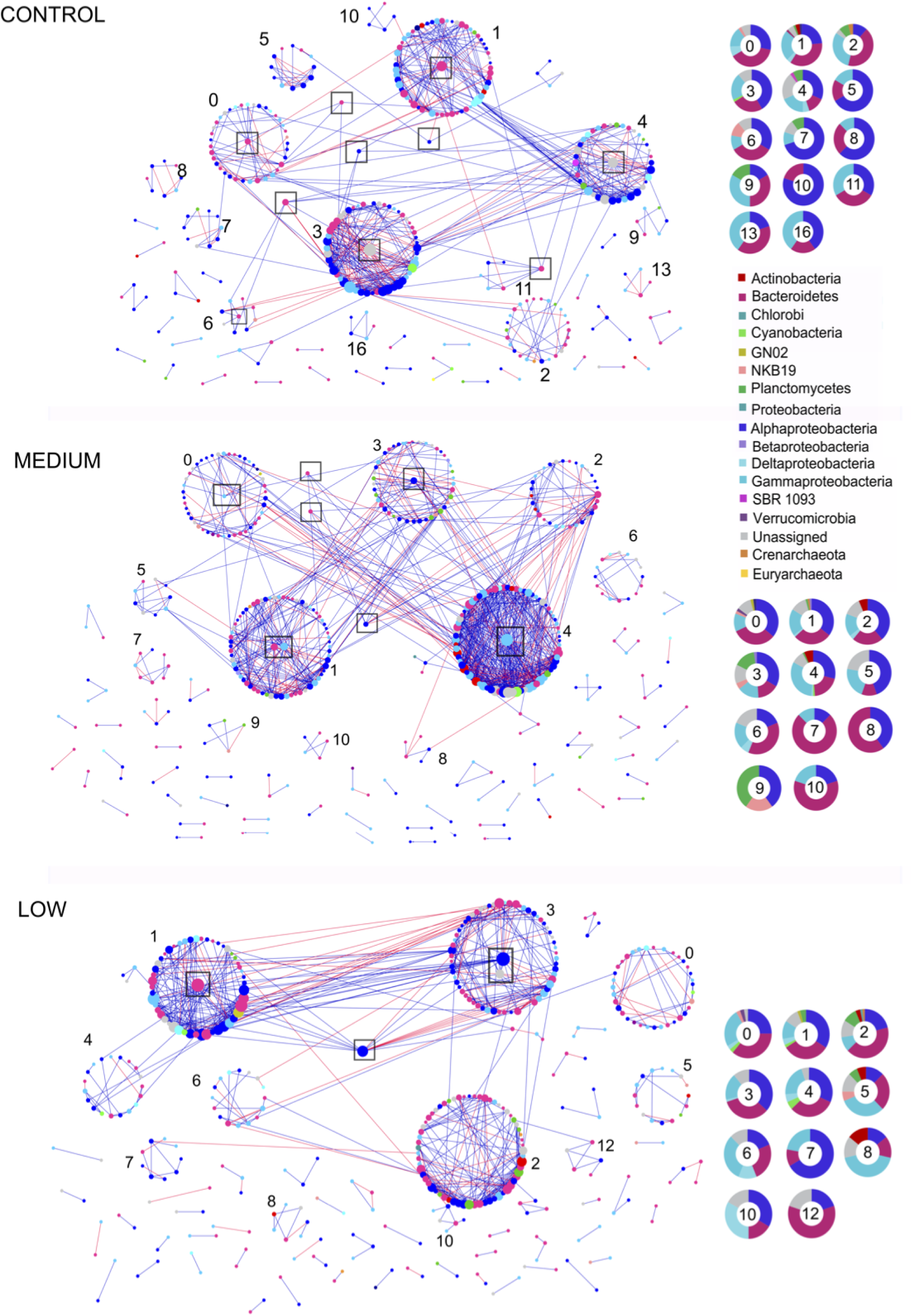
Ecological bacterioplankton association networks for the control, medium and low pH treatment groups. Nodes correspond to OTUs, where node size is plotted according to node degree, or connectedness. Node colours correspond to phyla and classes. Module taxa compositions are indicated with numbered donut charts, where numbers within a specific chart correspond to a particular module. Associations are defined by Spearman’s’ rho correlation coefficient and are deemed significant by random matrix theory (RMT). Positive co-occurrences are shown with a blue line whereas negative co-exclusions are indicated with a red line. Black boxes indicate within module keystones (module hubs) and between module keystones (connectors), respectively.

The number of modules in the medium and low networks were greater than the control network (Table S2). Modules were largest and most connected in the medium network, followed by the control and the low network, reflecting patterns observed from overall network structure. Intriguingly, the percentage of positive associations increased in response to acidification, where the low network had 77.82% positive associations, the medium network contained 77.25% and the control network only 74.70%, highlighting that reduced pH levels resulted in a slight increase in positive associations as opposed to co-exclusions (Fig. 4, Table S2). The overall taxonomic composition of the largest modules at both phylum and class level did not differ markedly between the three treatment groups, although some modules exhibited a dominance of different taxonomic groups (Fig. 4). Alphaproteobacteria, Bacteroidetes and Gammaproteobacteria were the dominant taxa in almost all modules and were also the dominant taxa in the overall bacterioplankton community. Actinobacteria occurred more frequently in the medium and low networks, and Euryarchaeota was found exclusively in the control network (Fig. S2 and Table S1). Unassigned sequences constituted a relatively high proportion in several modules, and were highest in the low network, suggesting that cryptic unknown species may increase in abundance in response to acidification in the SO. Furthermore, no modules consisted of a single class or phylum, showing a low degree of phylogenetic assortativity within the association networks, which suggest a potential high capacity for biotic interactions in SO microbial communities.

Network topological metrics can be computed for each node/OTU by assessing the within-module connectivity (*Zi*) as well as the between-module connectivity (*Pi*), which can indicate the potential ecological role of each OTU. These parameters can then be used to classify each OTU into four network-based categories; peripherals, connectors, module hubs and network hubs (Fig. 5a). Ecologically, peripherals may be considered as specialists, while connectors and module hubs may represent generalists and network hubs could be classified as super generalists. Additionally, the last three can be regarded as potential keystone taxa, on the sole basis of their high connectivity, since the overall network structure would be significantly altered if they were removed.

**Fig. 5:**
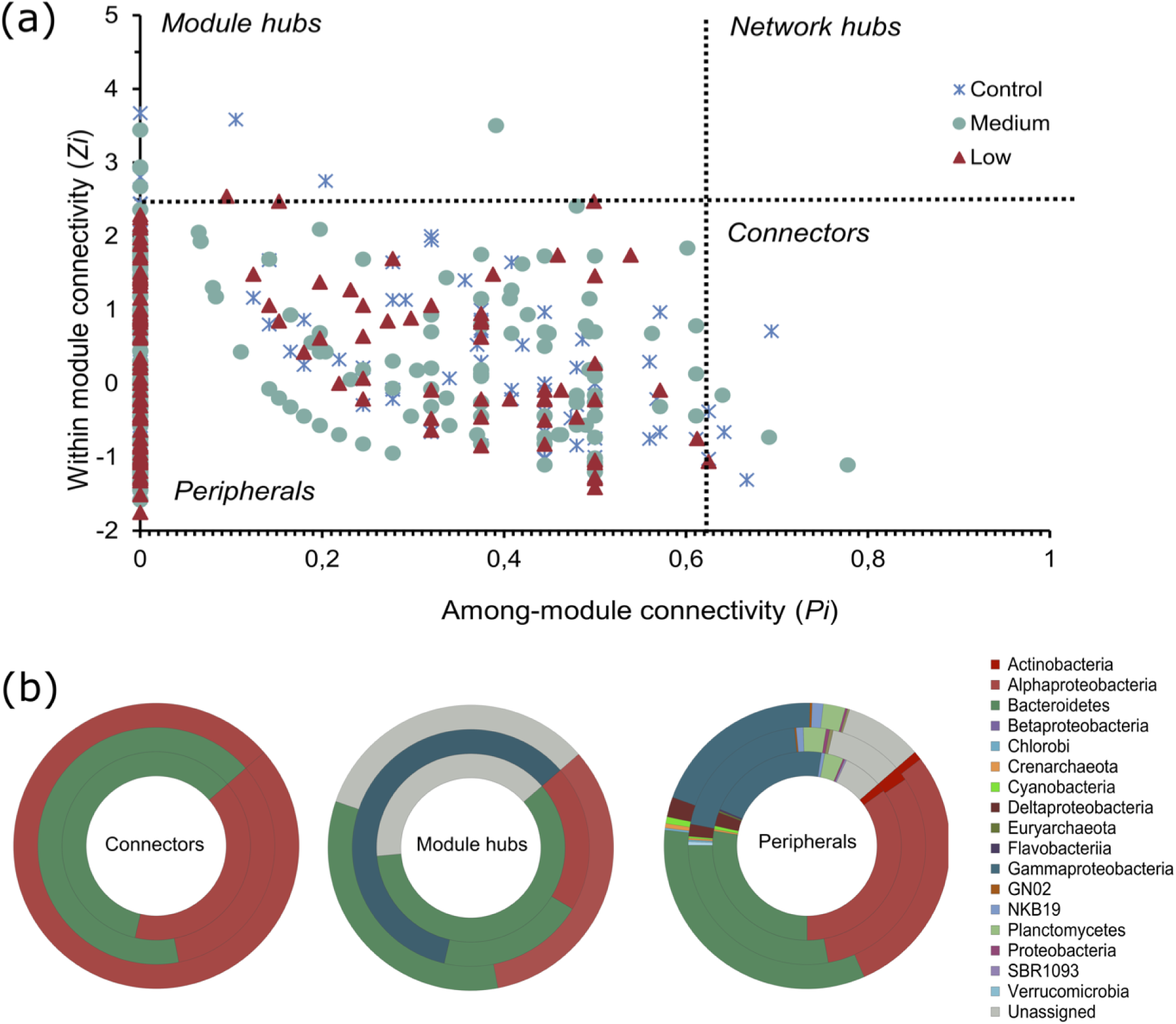
Plot of the within-module connectivity (*Zi*) and between-module connectivity (*Pi*) of each node or OTU in the three ecological networks and taxon relative abundances. Topological roles of OTUs are indicated either as peripherals (*Zi* ≤ 2.5, *Pi* ≤ 0.62), connectors (*Zi* ≤ 2.5, *Pi* > 0.62), module hubs (*Zi* > 2.5, *Pi* ≤ 0.62) and network hubs (*Zi* > 2.5, *Pi* > 0.62). The abundance of peripherals, connectors and module hubs between the three treatment groups is indicated in (b), where the control group is the centremost donut ring, the medium group the middle ring and the low group the outer donut ring.

The majority of OTUs in all three networks were classified as peripherals, with most edges occurring within their own modules (Fig. 5a). Several peripheral OTUs had a *Pi* value of 0, indicating that these nodes were connected solely to other OTUs within the same module (74% of OTUs in the control network, 70% in the medium network and 84% in the low). No network hubs (super generalists) were detected in any network. Interestingly, the relative abundance of several peripheral taxa changed in response to acidification (Fig. 5b, see Supplemental Information for details).

The number and taxonomic affiliation of module hubs and connectors detected (Table S3) differed noticeably between the three networks. Five module hubs were detected in the control network, five in the medium network and only three in the low network (Fig. 4, 5a). All control hubs consisted of phylum Bacteroidetes OTUs and unassigned sequences, with the orders *Flavobacteriales* and *Saprospirales*. Medium hubs belonged to classes Flavobacteriia, Gammaproteobacteria and Alphaproteobacteria. Low hubs consisted of Alphaproteobacteria, Flavobacteriia and unassigned bacteria. The relative abundance of module hubs ranged from 0.01% to 0.74%. The control network contained five connectors, the medium network contained three and the low network had only one. Control and medium connectors consisted of Alphaproteobacteria (order *Rhodobacterales* and *Rickettsiales*, respectively) and Flavobacteriia (order *Flavobacteriales*), whereas the low connector belonged to Bacteroidetes (order *Cytophagales*). Connector relative abundance ranged from 0.005% to 3.24%, with all but one OTU with an abundance of less than 1%, underlying the disconnection between species centrality and relative abundance. As hubs and connectors can be considered potential keystone taxa, Alphaproteobacteria and Flavobacteriia represented prominent putative keystone taxa within all networks, although differences between networks can be detected at finer taxonomic resolution (Table S3).

## DISCUSSION

### Diversity does not equal stability

Microbial community stability amidst perturbations is thought to be promoted by increased diversity, due to the link between diversity and ecosystem health and multifunctionality (Girvan et al., 2005; Delgado-Baquerizo et al., 2016; Shade, 2017). Similarly, loss of community diversity following environmental disturbance is often linked to community susceptibility. However, it has recently been suggested that assumptions of increased diversity leading to increased stability are false, at least for some ecosystems, as the role of diversity is dependent on the environment or ecosystem and disturbance itself (Shade, 2017). Community members are not necessarily interacting mutualistically in diverse compositions, and taxonomic diversity does not absolutely equate to functional diversity (Louca et al., 2016). In this study, we found that bacterioplankton alpha diversity is significantly higher in acidified compared to ambient mesocosms, in contrast to what has been hypothesised. Simultaneously, bacterioplankton community structure was significantly altered in response to CO_2_, with the relative abundances of several key community members significantly changing. We suggest that increased bacterioplankton diversity is an indication of community susceptibility, or “scramble” rather than stability in the Southern Ocean. The high diversity identified for bacterioplankton communities highlight that polar marine regions, and the SO specifically, contain high levels of unexplored diversity (Dickinson et al., 2016).

### Mycoplankton resistance to acidification

In contrast to our initial hypothesis, we did not observe a significant effect of ocean acidification on mycoplankton diversity. Instead, fungal community diversity and structure was stable, with the exception of Agaricomycetes, which increased in relative abundances at lower pH levels (Table S3). Agaricomycetes is a class of Basidiomycota that contains several yeast species (Jones, 2011; Jones et al., 2015), suggesting an increase in abundance of adapted fungal morphologies in response to acidification. However, the increase of Agaricomycetes phylotypes did not significantly affect overall fungal diversity or community structure. We argue that mycoplankton may contain a higher genetic plasticity and degree of physiological adaptation than bacterioplankton and are therefore resistant to future acidification. Culture dependant studies have shown that both terrestrial and marine fungi can rapidly adapt to a variety of environmental conditions, such as hot and cold temperatures and anoxia (Richards et al., 2012; Tsuji et al., 2013; Rédou et al., 2015). Additionally, fungi have larger pH tolerance ranges and optima for growth than bacteria (Rousk et al., 2010). Moreover, fungi possess chitin-rich cell walls (Taylor and Cunliffe, 2016), which may provide enhanced resistance. Source habitat significantly influences mycoplankton community structure (Panzer et al., 2015), further suggesting that SO fungal communities may be adapted to the pelagic environment and therefore less effected by pH variations. The lack of a significant mycoplankton community response in SO waters to pH levels predicted in the next 300 years, suggests the resistance of marine fungi to acidification in these systems.

### Altered organic matter cycling in a future acidic ocean

Microbially produced extracellular enzymes break down complex organic substrates, mediating heterotrophic carbon and nutrient recycling in marine environments (Arnosti, 2014). Enzymes measured for nutrient acquisition showed activity in all samples, indicating active bacterioplankton and/or mycoplankton heterotrophic communities within surface SO waters, since members of both assemblages are capable of producing each enzyme (Jones, 2011; Arnosti, 2014; Hoarfrost and Arnosti, 2017). The observed results of exoenzyme activities were congruent with our original hypothesis. Carbon and phosphorus degrading enzymes showed no substantial shifts in activity in response to pH treatment, suggesting constant, and not elevated carbon and phosphorus utilisation. Nitrogen acquiring enzymes, LAP and NAG, however, were negatively associated with pH, implying altered heterotrophic nitrogen recycling in a future more acidic ocean.

LAP and NAG activity responses contrasted to previous studies, which have found LAP activity to increase (Piontek et al., 2010; Endres et al., 2014) or remain unchanged (Yamada et al., 2008; Burrell et al., 2016). The decreased LAP and NAG activities suggest that nitrogen acquisition by bacterioplankton may be negatively impacted by pH, possibly due to increased physiological stress placed on microbial cells in a more acidic ocean. Bunse *et al*. (2016) found that bacterioplankton respond to a pH decrease of 0.2 units, applied over a period of only nine days, by increasing the expression of pH homeostasis genes. The exposure to lower extracellular pH likely increased physiological stress, resulting in intensified energy expenditure on cell maintenance, at the expense of bacterial growth (Bunse et al., 2016). Therefore, it is likely that the microbial communities producing LAP and NAG were negatively impacted by acidification. Reduced pH may have caused a loss of community members responsible for producing the nitrogen acquiring enzymes (Allison and Martiny, 2008a), ultimately decreasing cellular growth and fitness. Ultimately, microbial communities are essential produces of enzymes responsible for recycling organic matter, especially within nutrient deficient regions such as the SO. Although further research is necessary, alterations in bacterioplankton communities and their functions within the oligotrophic SO suggest negative future effects on ocean nutrient cycling.

### Bacterioplankton community network shifts as a response to acidification

Network topology, size and structure shifted markedly in response to reduced pH, with prominent differences between the control, medium and low bacterioplankton networks, congruent with our third hypothesis. Altered network structure and the loss of potential keystone taxa at lower pH levels suggest that marine microbial communities will be impacted by future acidification. Modules within networks may represent specific ecological niches (Olesen et al., 2007; Dupont and Olesen, 2012), and high interconnectedness between modules may indicate communication or co-operation between niches. The control group, although containing a large number of modules, did not contain a high number of connections between modules, and several between module connections were negative associations (Fig. 4). The large number of modules observed in the control group suggests that SO bacterioplankton communities form distinct niches and that both intermodular and intramodular predicted interactions may represent a balance between competition and mutualism. For example, microorganisms compete for nutrients and exhibit different growth constraints (Steele et al., 2011; Eiler et al., 2012; Brum et al., 2016) and also communicate via mechanisms such as quorum sensing (Hmelo, 2017). This study supports the notion that SO bacterioplankton communities are highly complex with a multitude of putative biotic interactions.

The higher network connectivity and complexity in the medium pH group suggests that bacterioplankton responded to a pH decrease of 0.4 units, associated with an increase in network stability (Fig. 4). Indeed, pH has been shown to alter community network structure (Barberan et al., 2012), resulting in larger and more complex phylogenetic and functional networks in response to elevated soil CO_2_ levels (Zhou et al., 2010; Zhou et al., 2011). Furthermore, network stability is thought to be promoted by increased modularity, as disturbances may spread more slowly through a modular than a non-modular network (Olesen et al., 2007). Module 4 contained the highest number of intra-module connections, where the majority of co-occurrences were positive (Fig. 4). The high intra-module connectivity is likely indicative of cooperation or mutualism between phylogenetically distant species within a specific niche. In contrast, several intermodular connections were negative, suggesting that increased physiological stress may result in more competitive interactions between niches at lower pH levels within the SO. This competition may be due to increased physiological stress on weaker community members, leading to the development of specialists with more efficient mechanisms for using the available nutrients. However, more research is needed to determine the cause of competition observed. Ultimately, future pH decreases of 0.4 units may promote increased bacterioplankton community interactions and network stability.

In contrast, the low network was the least complex, with reduced intermodular connectivity. This may suggest that microbial niches or guilds decreased the amount of interactions compared to the control and medium networks. This decreased cooperation may be due to the increased physiological stress placed on bacterioplankton communities at pH levels expected in the next 300 years. Acidification can promote direct physiological changes of microbial communities, whereby the increased energetic cost associated with cellular maintenance may lead to reduced microbial growth and fitness (Gilbert et al., 2008; Bunse et al., 2016). Decreased cellular fitness may result in altered functionality (Bunse et al., 2016) eventually perturbing community interactions. The increased stress induced by a pH decrease of 0.8 units likely resulted in the substantial alterations of the network structure, with potentially deleterious effects on bacterioplankton community structure in a future acidified ocean scenario.

### Keystone bacterioplankton in the SO and responses to acidification

Several potential keystone taxa were identified in all three networks, defined by their integral topological role in maintaining network structure and stability (Deng et al., 2012). The number of connectors decreased in the medium network, suggesting a switch between single microorganisms responsible for connecting niches to several community members linking different modules. The number of module hubs and connectors decreased dramatically in the low network. Loss of keystone members in a network decreases network stability and can lead to a collapse of network structure due to their high level of connectivity (Dunne et al., 2002; Olesen et al., 2007). The observed decrease in keystone nodes in the low network therefore correlates to the less stable network structure. Moreover, the reduced frequency of module hubs and connectors yet again suggests a lower frequency of community interactions or relationships in response to extreme levels of acidification.

The constructed networks are based on co-abundances and do not necessarily represent true biological interactions (Shi et al., 2016; Röttjers and Faust, 2019). However, co-occurrence networks have been instrumental for investigating the structure of microbial assemblages across biomes (de Menezes et al., 2015; Murdock and Juniper, 2019), in the detection of true viral host interactions (Chow et al., 2014; Soffer et al., 2015), the association of phylotype shifts to specific environmental parameters and functions (Valverde et al., 2015; Van Goethem et al., 2017), as well as in the detection of metabolic dependencies resulting in the culture of previously uncultivable microorganisms (Duran-Pinedo et al., 2011). Furthermore, networks provide valuable insight into complex community structures and stability, particularly in the face of perturbations, and give information on community organisation not portrayed by diversity metrics alone (Zhou et al., 2010). Thus, our results provide a framework for complex community predicted interactions in response to acidification (Fig. 6) where individual species interactions may be experimentally tested in future studies.

**Fig. 6:**
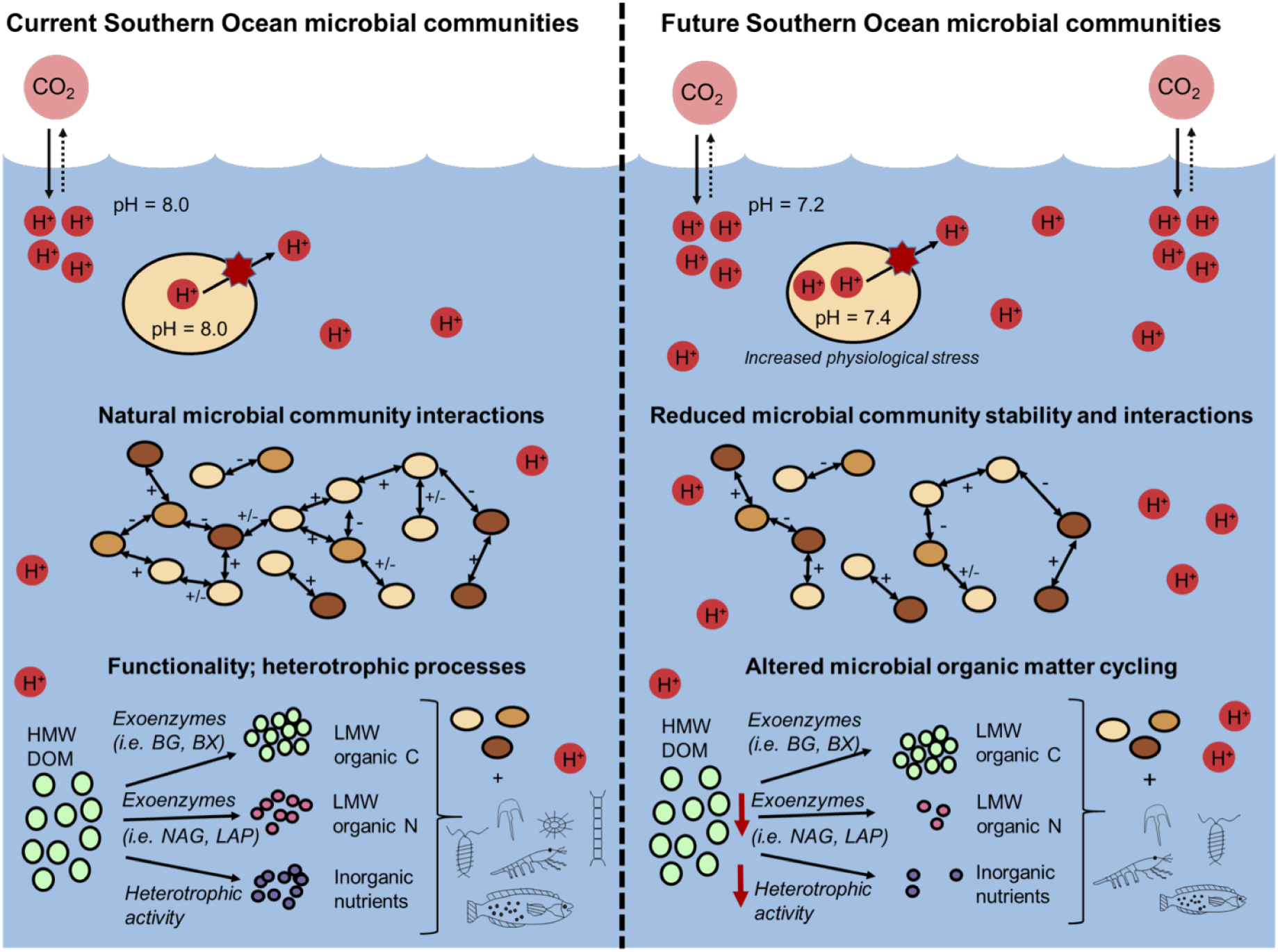
Conceptual diagram depicting ocean acidification effects on SO surface microbial community structure and functionality. The left and right panel indicate current and future pH levels, respectively. Ovals represent microbial communities, where the lightest colour is indicative of bacteria, the middle colour archaea and the darkest colour fungi. Headings indicate different processes; the first panel shows the process of CO_2_ exchange with the atmosphere and the associated consequences on ocean H^+^ concentration, pH and a bacterial cell (bacterial cell = yellow oval; proton pump = red star); the second panel indicates current and future microbial community interactions; the last panel shows functional processes utilised to degrade high molecular weight DOM, the subsequent products, and the usage of the resultant products by microbial communities and the higher trophic system. Abbreviations are as follows: HMW = High molecular weight; LMW = Low molecular weight; C = carbon; N = nitrogen; H+ = hydrogen ions; BG = beta-glucosidase; BX = beta-xylosidase; NAG = N-acetyl-glucosaminidase; LAP = leucine aminopeptidase.

In summary, we have shown that ocean acidification significantly alters bacterioplankton, but not mycoplankton, communities. Exoenzyme activities involved in nitrogen acquisition were substantially reduced at lower pH levels, suggesting a diminished availability of nutrients in surface waters and amended nutrient recycling. Additionally, bacterioplankton network structure and keystone taxa were significantly impacted by increased CO_2_. We propose that future acidification may have adverse consequences on microbial communities and biogeochemical cycling, particularly within regions most sensitive to future climatic changes.

## MATERIALS AND METHODS

### Sampling and mesocosm experiment

Seawater was collected during the Marion Island Relief Cruise aboard the RV SA Agulhas II from April 2016 to May 2016. Approximately 500 L of seawater was collected from surface waters (5 m) using the underway system at a single site (46°00.065’ S, 38°00.37’ E). The water was distributed into nine mesocosms (50 L volume) with the sterile drums filled to near capacity to keep the headspace small (volume 5 – 10ml) in order to minimize the exchange of CO_2_. The mesocosms consisted of three biological replicates for each pH treatment group, where treatments are defined as follows: a control (i.e. pH 8.0) which is the natural pH of the environment, a medium pH, which mimics predicted pH decreases within the next 100 years (0.4 units, i.e. pH 7.6), as well as a low pH treatment which exceeds the predicted decrease in pH within the next millennium (i.e. pH 7.2). All values were allowed to deviate by ± 0.05 pH units in order to account for natural variability within the SO environment (Kapsenberg et al., 2015). The mesocosms were exposed to light on a 12 hr day/night cycle, where light was allowed to penetrate the top of the mesocosms in order to mimic the natural light conditions at 5 m. Mixing of mesocosms was performed via consistent O_2_ bubbling (HAILEA Air Compressor, Guangdong, China), as well as manual stirring throughout the experiment. Each treated mesocosm contained a pH probe (MODEL PH-301C, SAGA, Tainan City, Taiwan, pH accuracy ± 0.1% + 2 digits) which consistently recorded the seawater pH. Physico-chemical variables within each mesocosm such as temperature, salinity, conductivity and pH, were also manually recorded twice a day throughout the duration of the experiment using a HANNA probe (MODEL HI 98195, Padova, Italy, pH accuracy ± 0.02 units, temperature accuracy ± 0.15°C, conductivity and salinity accuracy ± 1% of reading). Changes in pH were induced via CO_2_ bubbling and the experiment was run for a total of 21 days, during which pH changes were induced on the first day (day 0) of the experiment. Both O_2_ and CO_2_ bubbling were introduced into mesocosms via sterile plastic tubing, where the amount of gas was controlled using a regulator. Samples were collected every 7 days at noon, including at the start (day 0) of the experiment. Physico-chemical variables such as temperature, salinity, and nutrients were recorded throughout the experiment (see Fig S1).

### Enzymatic activities

Seawater collected from each sampling point in the mesocosm experiment was assayed for the activity of 5 enzymes, including β-glucosidase (BG), β-xylosidase (BX), β-N-acetyl-glucosaminidase (NAG), alkaline phosphatase (AP), and leucine aminopeptidase (LAP). The potential activities of these enzymes were tested using fluorescent assays, and enzyme activities were performed and calculated as described previously (Sinsabaugh, 1994; Papanikolaou et al., 2010; Hoarfrost and Arnosti, 2017). Briefly, 200 µL of seawater was added to 96-well black flat-bottom microplates (Greiner Bio One, Frickenhausen, Germany), where four replicate wells were used per sample. A concentration of between 1 mM to 1.5 mM of each enzyme substrate was then added to the 96 well plates. Plates were incubated for 2 hours at 4°C in the dark. Fluorescence was measured from the black 96 well plates using a Spectramax® Paradigm Multi-Mode Microplate Reader (Molecular Devices, USA) with an emission wavelength of 450 nm and an excitation wavelength of 360 nm.

### Molecular analysis

Seawater (2 L) collected from each sampling point in the mesocosm experiment was filtered through 0.2 µM pore cellulose acetate membrane filter (Sartorius Stedim Biotech, Göttingen, Germany). Nucleic acid was subsequently extracted from each of the filtered samples using the PowerWater DNA Isolation Kit (MoBio, Carlsbad, USA), according to the manufacturer’s instructions. DNA was sent to MR DNA (www.mrdnalab.com, Shallowater, USA) for paired-end sequencing (2 × 300bp) of the 16S rRNA gene and the internal transcribed spacer (ITS) 1 rDNA variable region on the Illumina MiSeq platform (Illumina, San Diego, CA, USA), as detailed previously (Phoma et al., 2018; Vargas-Gastélum et al., 2019). The marine specific primer pair 515F-Y and 926R (Walters et al., 2016) was used to PCR amplify hypervariable regions V4 and V5 of the bacterial and archaeal 16S rRNA gene. ITS region 1 was targeted and PCR amplified using the primer pair ITS1F (Gardes and Bruns, 1993) and ITS2 (White et al., 1990). All samples were barcoded on the forward primer to allow for multiplex sequencing.

Sequences were analysed using QIIME v 1.8.0 (Caporaso et al., 2010). Sequences were demultiplexed according to barcode identity and were quality filtered according to QIIME default parameters, with exceptions for the ITS dataset. For the ITS dataset, the p score applied was altered to 0.45 to ensure that all reads >200bp were included in the dataset, provided that they met the other default quality parameters (i.e. q 19, r 3, etc.). Chimeras were detected both *de novo* and against the Greengenes reference library (DeSantis et al., 2006) for 16S rRNA genes and the UNITE reference database for the ITS region (Kõljalg et al., 2013) using usearch61 (Edgar et al., 2011). Operational taxonomic units (OTUs) were clustered at a similarity threshold of 97% using the uclust method and the Greengenes and UNITE reference database for 16S rRNA gene and ITS region datasets, respectively.

16S rRNA gene OTUs were assigned a taxonomy using the Greengenes (DeSantis et al., 2006) reference database. Taxonomy was assigned to ITS region OTUs using a combination of the UNITE database in QIIME and the Warcup Fungal ITS training set in the RDP classifier (Wang et al., 2007; Kõljalg et al., 2013; Deshpande et al., 2016). For the RDP classifier, taxonomic classifications were kept at the default confidence threshold of equal to or greater than 80% for classification to the root only, after which the default confidence threshold was decreased to 50% for all subsequent assignments. Following taxonomy assignment, singletons were removed from the datasets and both datasets were rarefied to the lowest number of sequences per sample (53,267 for 16S rRNA gene and 10,370 for ITS region), using a single rarefaction. All sequences are available on the SRA database under the following accession number: SRP126760

### Network analysis

Ecological association networks were constructed using Random Matrix Theory (RMT) through the Molecular Ecological Network Analysis (MENA) pipeline (http://ieg2.ou.edu/MENA/) (Deng et al., 2012). Individual networks were constructed for the three treatment groups (i.e. one low, medium, control) where all days were combined, excluding day zero (i.e. nine biological replicates per network, see Supporting Information for details). All networks were visualised in Cytoscape v 3.4.0 (Shannon et al., 2003).

### Statistical analysis

All statistical analyses were performed using R statistical software version 3.2.2. (R Core Team, 2013). Alpha diversity was calculated using the *vegan* package (Oksanen et al., 2011). Statistically significant differences in diversity measures between each treatment group were determined via the *aov* function (analysis of variance, ANOVA) in *vegan* and the post-hoc TukeyHSD test. Microbial community structure was visualised by non-metric multidimensional scaling (nMDS) ordination plots with the *metamds* function of *vegan* using Bray-Curtis dissimilarity and visualised using *ggplot2*. Significant differences in microbial community structure were determined via PERMANOVA (permutational analysis of variance) using the *adonis* and *betadisper* functions in *vegan*. The *aov* and *tukeyHSD* functions of *vegan* were used to determine significant differences between treatment groups for each of the enzyme activities. Correlations between taxon abundances, enzyme activities and pH were calculated using Spearman’s rho, via the *rcorr* function of the *Hmisc* package. Relationships between pH and enzyme activities, bacterial, archaeal and fungal diversity and taxon abundance were modelled using linear regressions (*lm* function).

## ACKNOWLEDGEMENTS

We are grateful to the National Research Foundation (NRF) (Grant ID 100052 SZdS) the South African National Antarctic Programme (SANAP 110717), and the University of Pretoria for funding. TPM also wishes to acknowledge the Fulbright Visiting Scholar Program for providing sabbatical funding. We wish to acknowledge Mr. A. van der Walt and Mr. M. Clayton for useful discussions on the content of the manuscript.

